# Identification of the siderophore schizokinen and its derivatives by LCHRMS and mass-tandem fragmentation

**DOI:** 10.64898/2026.05.05.723046

**Authors:** Ignacio Sottorff

## Abstract

Biological metal chelators are of great interest for investigation due to their capacity to retain or mobilize metals from the environment. While some biological and bioinspired chelators find use in medical applications, others are promising platforms for the mining or recycling of technologically important metal ions. In particular, the siderophores, which are primarily iron chelators, have been studied. Four siderophores of relevance are schizokinen and its derivatives, which have been isolated from bacterial and algae cultures, in addition to soil. These siderophores have shown metal chelating activity with different metals such as iron, copper, and aluminum. In the time of metabolomics, it is required to unambiguously determine the identity of the produced siderophores as quickly as possible. Thus, Liquid Chromatography coupled to High Resolution Mass Spectrometry and mass-tandem fragmentation (LC-HRMS-MS) provides a quick and applicable alternative for identification of schizokinen and its derivatives. Here, we report an analytical method for the identification and potential quantification of the schizokinen siderophore series. We developed a working method through LC-HRMS-MS, which provides the unequivocal identification of the four schizokinen derivatives, which has not been reported to date. Additionally, we constructed the molecular network for the four molecules to enable their identification using the Global Natural Products Social Molecular Networking (GNPS) platform. Most importantly, this contribution can help speed up the characterization of schizokinen producers and facilitate the dereplication process of siderophores.

## Introduction

Metal chelators have become relevant because of their property to bind iron and other metals (Schalk et al. 2011). The metal binding feature can find applications in metal recycling (Martinez-Gomez et al. 2016), mining (Williamson et al. 2021), as well as in medicine (Swayambhu et al. 2021). It has also been shown that metal chelators can provide benefits for increasing crop yields (Ansari et al. 2017), which is needed for the stability of the global food supply chain. The most prominent group of metal binding molecules are the low molecular weight, high-affinity Fe(III)-chelators, also known as siderophores. To date, there are 5 classes of siderophores (**Fig 1**): hydroxamates such as desferrichrome (Emery and Neilands 1961), catecholates such as enterobactin (Raymond et al. 2003), α-hydroxy-carboxylates such as rhizoferrin (Drechsel et al. 1991), diazeniumdiolates such as gramibactin (Hermenau et al. 2018), and hybrids bearing multiple functional groups such as pacifibactin (Hardy and Butler 2019). Siderophores have also been known to be a key player in the bacterial infection process as the signal that regulates virulence factors to produce the disease (Bonneau et al. 2020; Holden and Bachman 2015; Kramer et al. 2020; Lamont et al. 2002).

**Figure 1.**
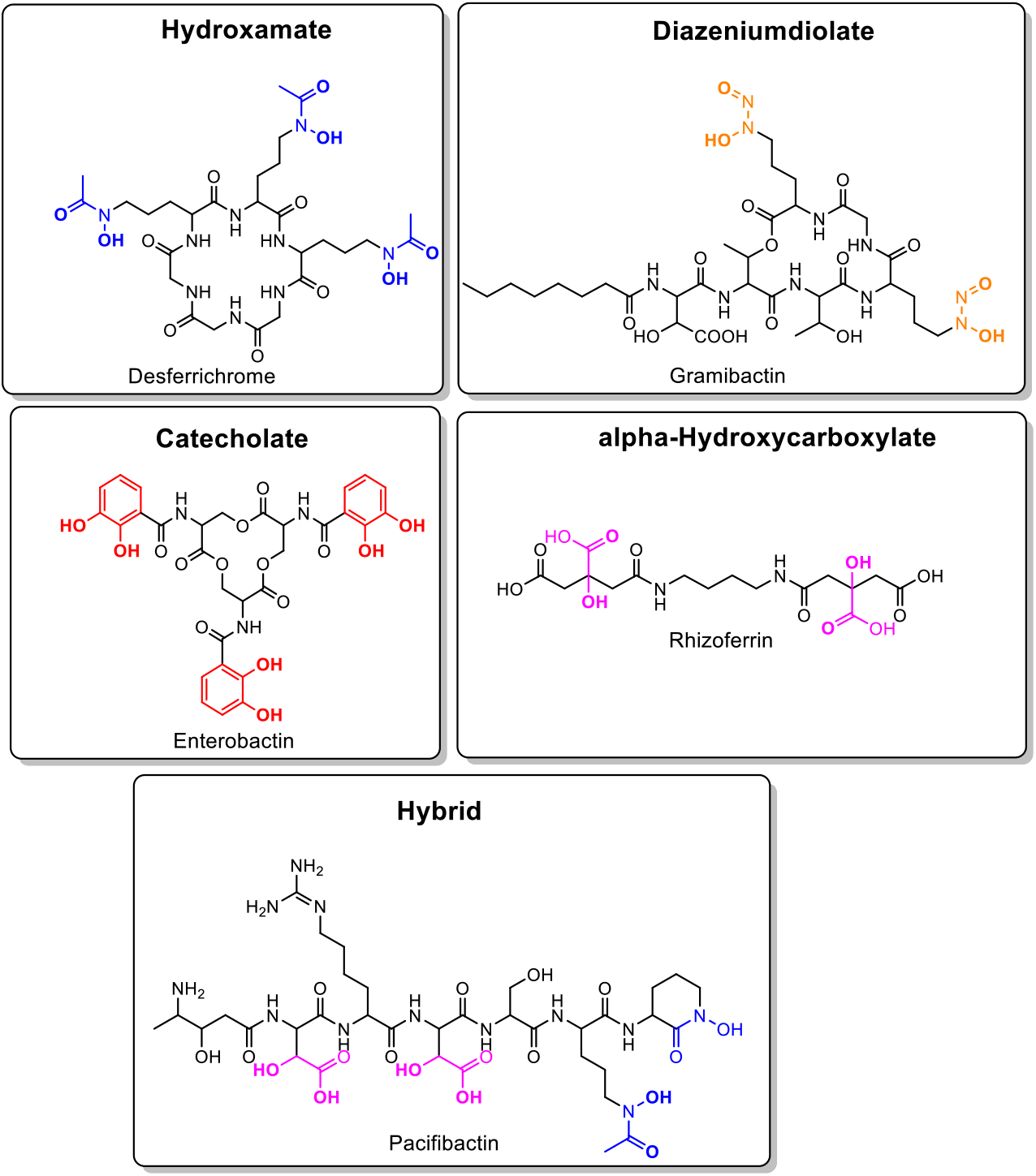
Siderophore classes. Functional groups that lead to their classification in color and coordinating donor atoms in bold.

Usually, siderophores are not larger than 2000 Da, which facilitates their unambiguous detection by liquid chromatography coupled with mass spectrometry. Tandem mass spectrometry and the fragmentation of a single molecule into distinctive fragments yields a well-defined fingerprint that can be used for the unequivocal identification and quantification of the analyte of interest (McLafferty 1981). Thus, the availability of a method which combines three layers of information for the identification of siderophores (retention time, exact mass, and mass fragmentation patterns), can contribute enormously to siderophore detection. Moreover, the development of public mass tandem fragmentation platforms (Allen et al. 2015; Allen et al. 2014), which allow the *in-silico* prediction and confirmation of the mass-spectrometry fingerprint produced by a given molecule, has facilitated the assignment of the most common fragment ions found in mass tandem experiments. In addition, new tools for the analysis of MS2 data have been generated, such as the Global Natural Products Social Molecular Networking (GNPS)(Yang et al. 2013). The GNPS allows the generation and visualization of molecular networks from the MS2 data, which facilitate and accelerate the characterization of known metabolites, as well as the targeting of unknown analogues that bear similar fragmentation patterns to identified molecules.

Schizokinen, a siderophore composed by one citric acid unit which is symmetrically linked with two residues of 1-amino-3*-*(*N*-hydroxy-*N*-acetyl)-aminopropane, was the first molecule of the schizokinen series to be characterized (Mullis et al. 1971). Subsequently, three further derivatives were identified: schizokinen A, N-deoxy-schizokinen and N-deoxy-schizokinen A (Hu and Boyer 1995; Mullis et al. 1971), which are shown in **Fig 2**. Thus far, schizokinen and its derivatives have shown to be chelators of iron (Mullis et al. 1971), copper (Arceneaux et al. 1984; Clarke et al. 1987), and aluminum (Hu and Boyer 1996). Schizokinen has shown growth promotion activity in bacterial cultures (Byers et al. 1967). It has been proposed that the production of the two schizokinen cyclic imide derivatives, schizokinen A and N-deoxy-schizokinen A, are an artifact that occur under acidic conditions, which produces the cyclization of schizokinen and N-deoxy-schizokinen to form the cyclized derivatives (Hu and Boyer 1995). Similar observations have been made upon heating schizokinen to transform it into its cyclic derivative, schizokinen A (Mullis et al. 1971). For schizokinen, it has been reported that the two hydroxamates in conjunction with the hydroxyl and carboxyl groups of the citric acid backbone generate the needed six coordination sites required for iron chelation (Plowman et al. 1984). Up to now, schizokinen and its derivatives have been isolated from bacterial (*Bacillus megaterium* and *Rhizobium leguminosarum*) and algae (*Anabaena sp*.) cultures (Hu and Boyer 1995; Simpson and Neilands 1976; Storey et al. 2006), as well as from rice fields soil (Akers 1983).

**Figure 2.**
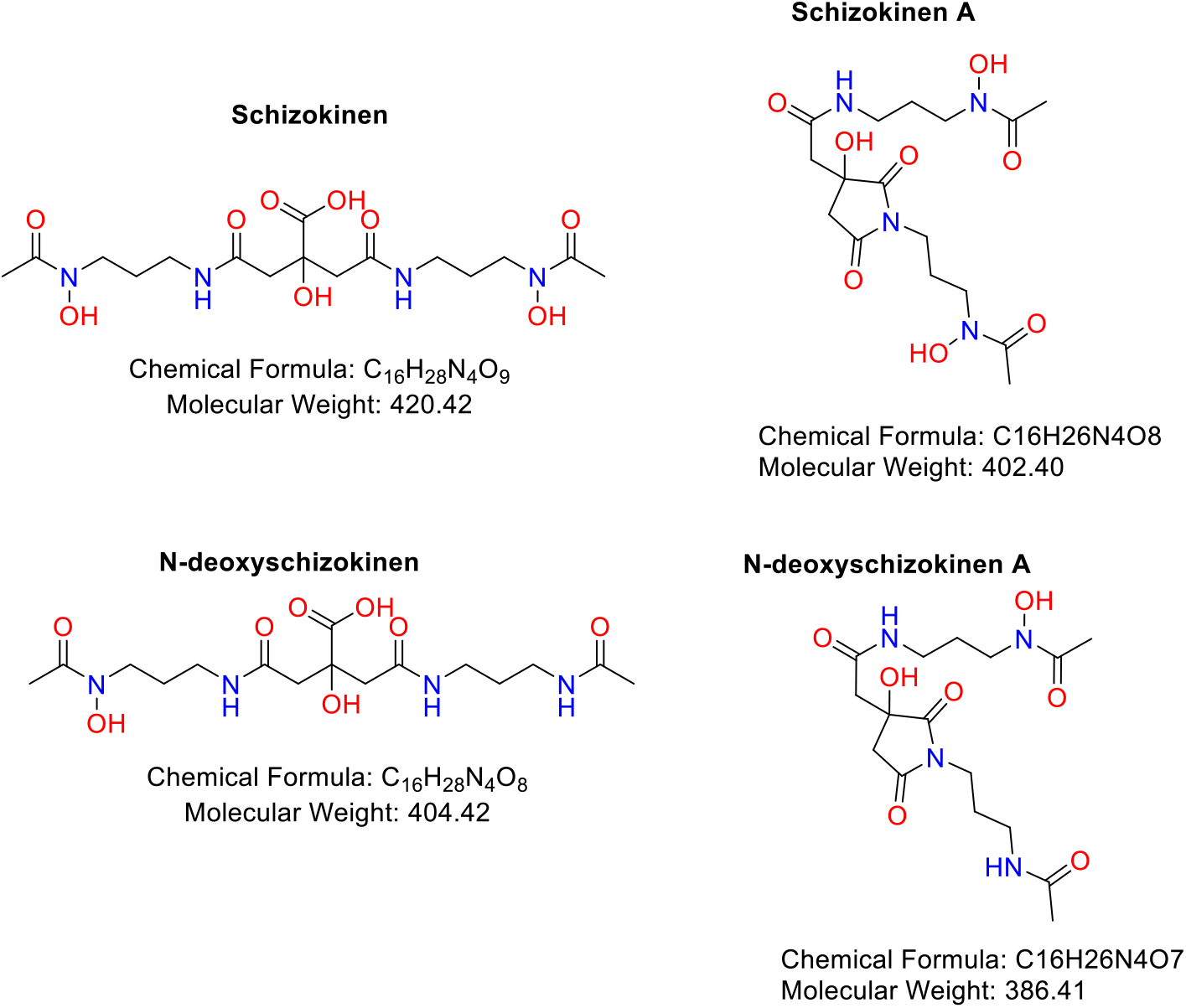
Schizokinen and its derivatives.

The open forms of schizokinen (schizokinen and N-deoxyschizokinen) have two available chelating functionalities, hydroxamate and α-hydroxycarboxylate. In contrast, the cyclized forms (schizokinen A and N-deoxyschizokinen A) have the known hydroxamate functionality to chelate metals. The iron coordination properties of schizokinen (pFe^III^_[pH=7.4]_=26.8) and its imide derivative (pFe^III^_[pH=7.4]_=27.8) have been studied, revealing that they have a higher affinity for iron(III) than desferrioxamine B (pFe^III^_[pH=7.4]_=25.0) (Chuljerm et al. 2019). Ultimately, schizokinen and its derivatives have been evaluated as ^68^Ga-PET agents for diagnostic applications (Chuljerm et al. 2020).

We report, herein, a methodology for the identification and characterization of the schizokinen series using liquid chromatography coupled to mass and mass tandem spectrometry, as well the molecular network of schizokinen and its derivatives. The implementation of this methodology can help to unequivocally determine the production and presence of schizokinen and its derivatives in biological, clinical and soil samples.

## Materials and Methods

### Siderophore standard

Schizokinen, deoxyschizokinen, schizokinen A and deoxyschizokinen A were obtained from Biophore Research Products (Tübingen University, Tübingen, Germany) as a single mix of schizokinen derivatives and stored dry at -20°C. Schizokinen and its derivatives were dissolved in methanol and water (1:1). The injection concentration and volume for the molecule mixture was 0.1 mg/mL and 50 µL. The samples were filtered through PTFE 0.22 µm filter (Fisher scientific, China) prior to their injection. After the injection, the samples were stored at -20°C.

### HPLC methodology

For the analysis of schizokinen and its derivatives a method was developed using a 30-minute gradient of water and acetonitrile supplemented with 0.1% LCMS grade formic acid (Fisher chemical, Geel, Belgium), and employing an Agilent 1260 Infinity II HPLC system (Agilent Technologies Inc., Santa Clara, California, USA). The chromatographic columns used were NUCLEODUR C18 Isis, 5 µm, 250×4.6 mm from Macherey−Nagel (Dueren, Germany) and Reprosil Gold 120 C18, 5 µm, 250×4.6 mm from Dr. Maisch (Ammerbuch-Entringen, Germany). The gradient used in both columns was as follows: 0 min 90% A 10% B, 2 min 90% A 10% B, 20 min 0% A 100% B, 23 min 0% A 100% B, 28 min 90% A 10% B, 30 min 90% A 10% B, where A is LCMS grade water (Merck, Darmstadt, Germany) and B: LCMS grade acetonitrile (VWR, Fontenay-sous-Bois, France) both supplemented with 0.1% LCMS grade formic acid (Fisher chemical, Geel, Belgium). The flow rate used was 0.7 mL/min. The instrument was controlled through MassHunter (Agilent Technologies Inc., Santa Clara, California, USA).

### Detection

The detection of the molecules was performed by two means. Using the diode array detector (DAD) at 220 nm of an Agilent 1260 Infinity II HPLC system (Agilent Technologies Inc., Santa Clara, California, USA) and total ion current (TIC) chromatogram (including the extracted-ion chromatogram, EIC) by using an Agilent 6530 Q-TOF LC/MS mass spectrometer (Agilent Technologies Inc., Santa Clara, California, USA).

### Mass spectrometry and mass-tandem fragmentation

For the determination of the exact mass of the molecules an Agilent 6530 Q-TOF LC/MS mass spectrometer was used, equipped with a Jet Stream Technology Ion Source (Agilent Technologies Inc., Santa Clara, California, USA). The parameters applied for the determination of the exact mass were as following: positive mode, gas temperature: 350°C, drying gas flow: 10 L/min, nebulizer pressure: 45 psi, sheath gas temperature: 400°C, sheath gas flow: 12L/min, VCap: 3500 V, nozzle voltage: 0 V, fragmentor: 100 V, skimmer: 65 V, oct RD Vpp: 750 V. The mass range used for the experiments was 50-1000 Da. The same parameters were used for the mass-tandem fragmentation experiments, but this time using the targeted MS/MS function. The collision energies for all the molecules, but N-deoxy schizokinen A were 10, 20 and 30 eV (schizokinen, schizokinen A and N-deoxy schizokinen), while that for N-deoxy schizokinen A, 10, 15 and 20 eV were used. The instrument was controlled through MassHunter (Agilent Technologies Inc., Santa Clara, California, USA). The same software was used to process the data.

### Mass-tandem fragments prediction

To compare and assign the obtained mass-tandem fragmentation pattern, the online platform CFM-ID was used (Allen et al. 2014). CFM-ID is a web server which can predict the most likely tandem mass spectra (MS/MS) fragments of a given molecule. The web server uses an algorithm based on competitive fragmentation modelling, which has been trained with machine learning techniques (Allen et al. 2015). The platform is available at http://cfmid.wishartlab.com.

### Molecular networking

To construct the molecular networking for the schizokinen series, we followed the procedure of the GNPS platform (Yang et al. 2013). Each molecule was fragmentated with three different collision energies, 10, 20 and 30 eV using an untargeted MS/MS method in an Agilent 6530 Q-TOF LC/MS mass spectrometer. The parameters applied for the data acquisition were as following: positive mode, gas temperature: 350°C, drying gas flow: 10 L/min, nebulizer pressure: 45 psi, sheath gas temperature: 400°C, sheath gas flow: 12L/min, VCap: 3500 V, nozzle voltage: 0 V, fragmentor: 100 V, skimmer: 65 V, oct RD Vpp: 750 V. The mass range used for the experiments was 50-1000 Da. Next, the data was analyzed and curated for accuracy. Subsequently, the data was submitted to the GNPS platform, and the network constructed. The visualization of the network was done using Cytoscape (Version: 3.8.2).

## Results and Discussion

### Standard resuspension

A 1:1 mixture of water/methanol was found most suitable to dissolve the standard, as neither pure methanol nor acetonitrile produced clear solutions at the desired concentration (0.1 mg/mL). Other studies have performed a similar strategy but using 1:1 water/acetonitrile instead of 1:1 water/methanol (Storey et al. 2006). In the case that crude extracts from different sources (bacterial, fungal, or environmental) need to be analyzed for schizokinen and its derivatives, we suggest dissolving the extracts in the same mix than this report, methanol/water (1:1), because it can help to reduce the extract complexity, work as a clean-up step and dissolve most if not all the schizokinen derivatives.

### HPLC method development and detection

The development of the HPLC method for the whole schizokinen series was firstly developed using a gradient of pure acetonitrile and water on a NUCLEODUR C18 Isis column from Macherey-Nagel. However, this first methodology was not successful in producing the separation of the four molecules, because schizokinen and N-deoxy schizokinen had the same retention time: 4.2 min (not shown). By the supplementation of 0.1% formic acid to the mobile phase, we were able to achieve the separation of the four analytes (**Fig 1A-E**) in a single injection. Subsequently, we tested a second column to determine if the developed method can also work in a different vendor C18 column. As it can be observed in **Fig 3** and **4**, a different C18 column with identical physical dimensions produced a similar result to our primary chromatographic separation results, confirming that the present chromatographic method is not column specific and thus, robust.

**Figure 3.**
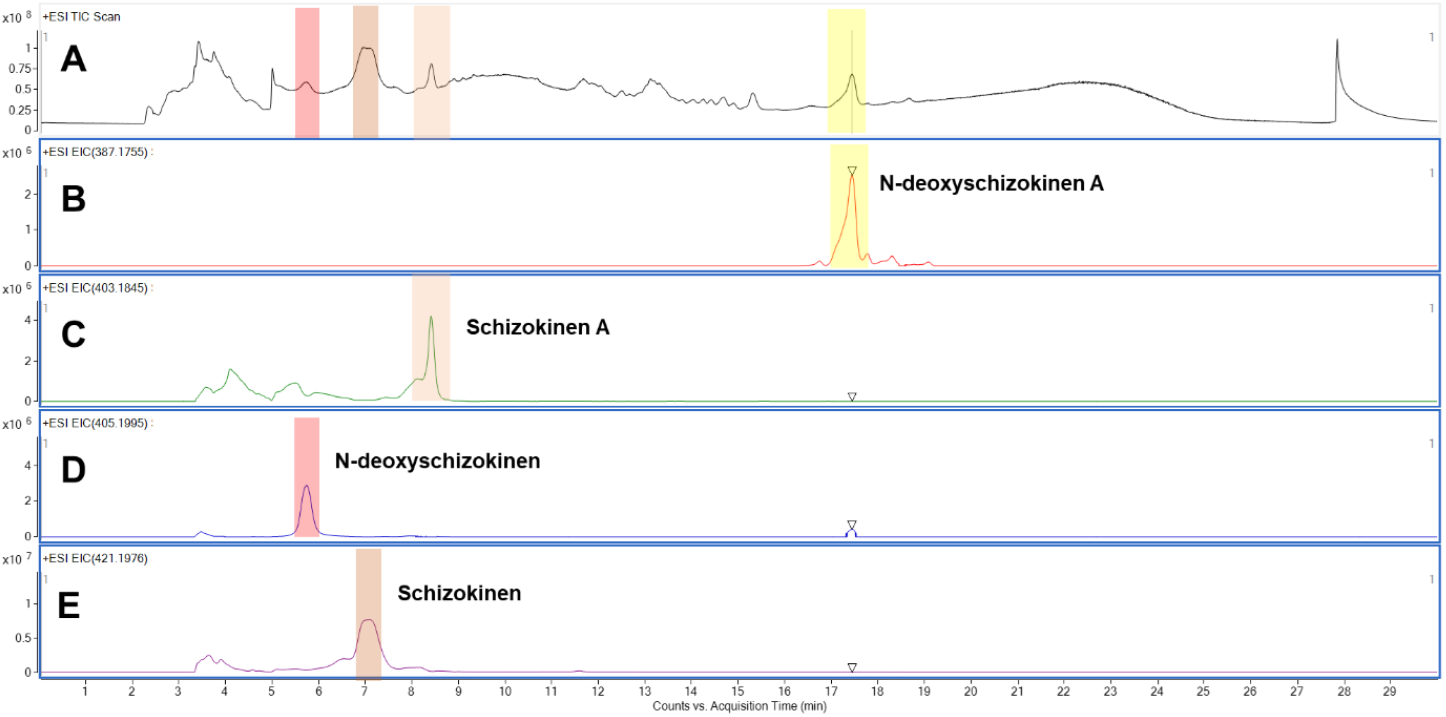
Chromatogram of the schizokinen series standard performed on a NUCLEODUR C18 Isis, 5 µm, 250×4.6 mm from Macherey-Nagel (Dueren, Germany). Total ionization chromatogram (TIC) for the schizokinen series (3A) and the extracted ion chromatogram (EIC) for each derivative (3B-D).

Previous studies have developed HPLC methods using isocratic and gradient methodologies in combination with C18 columns, (Hu and Boyer 1995; Plowman et al. 1984).However, these studies did not report whether the used HPLC methodologies had good resolution and separation for the whole set of schizokinen derivatives.

The detection of the desferri-schizokinen series by DAD is challenging, due to the lack of an appropriate chromophore. Nevertheless, at 220 nm we were able to observe a weak signal that may help for the primary identification of the molecule, as well as the chemical purification of the schizokinen series. The same wavelength has been used previously for the purification of schizokinen (Hu and Boyer 1995). Nevertheless, upon formic acid supplementation of the mobile phase, any visible response at 220 nm is lost, so that another method of detection was needed. Therefore, we used the total ion current (TIC) and extracted-ion chromatogram (EIC) signal of the mass spectrometer as a reliable mean of detection of the analytes. Alternatively, an evaporative light scattering detector (ELSD) might be suitable for the schizokinen series as well.

### Exact mass and retention determination

After the LCMS runs, we firstly analyzed the data and searched for the targeted molecules using the total ion current (**Fig 3A and 4A**). Subsequently, we obtained the individual extracted-ion-chromatogram (EIC) for each of the molecules, as shown in **Fig 3B-E** and **Fig 4B-E**. As it can be seen in **Fig 3** and **4**, the HPLC methodology produced a satisfactory separation for all four target molecules in two different vendors C18 columns, yielding the following retention times for the molecules in order of elution. For the NUCLEODUR C18 Isis column: 5.73 min for N-deoxyschizokinen (**3D**), 7.05 min for schizokinen (**3E**), 8.41 min for schizokinen A (**3C**) and 17.4 min for N-deoxyschizokinen (**3B**). For the Reprosil Gold column: 6.09 min for N-deoxyschizokinen (**4D**), 7.52 min for schizokinen (**4E**), 8.99 min for schizokinen A (**4C**) and 18.25 min for N-deoxyschizokinen (**3B**).

**Figure 4.**
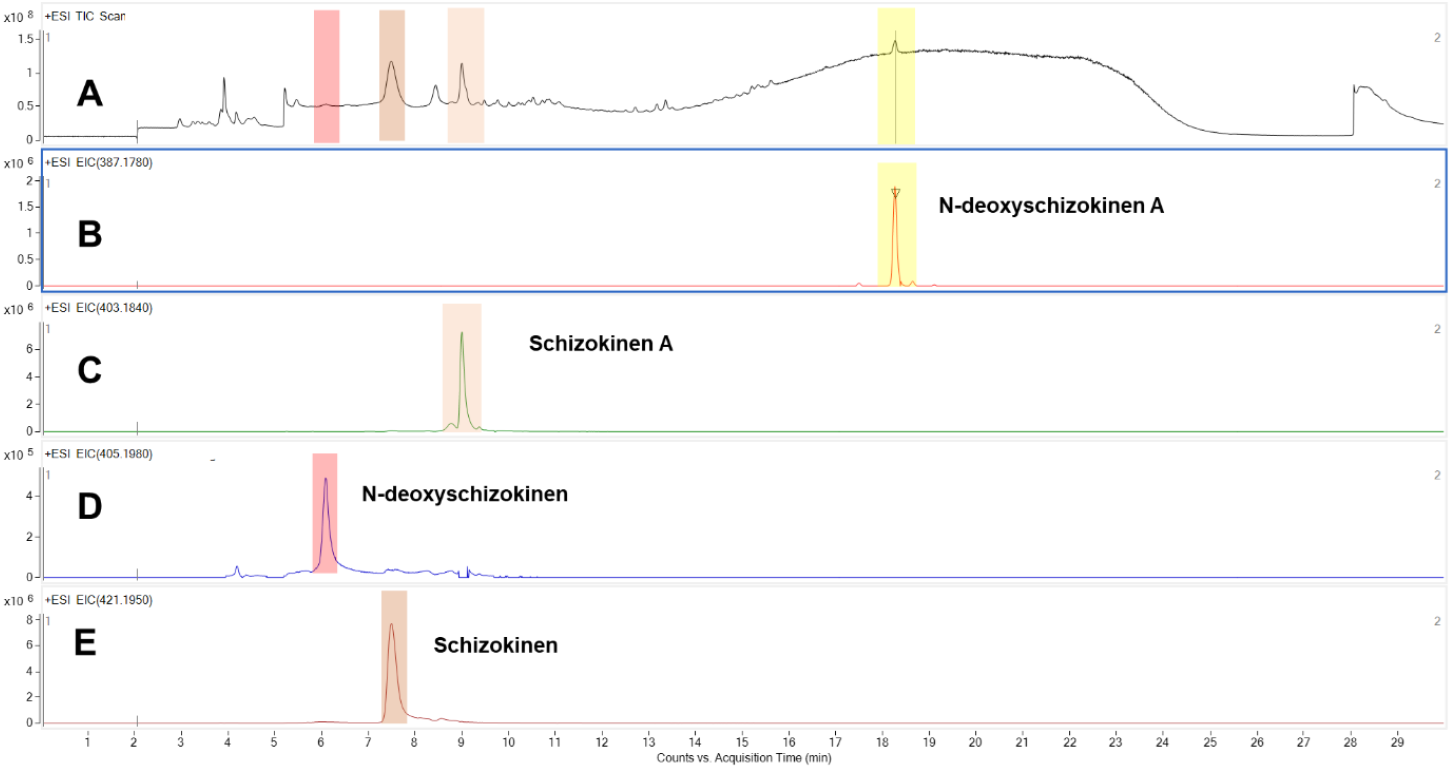
Chromatogram of the schizokinen series standard performed on a Reprosil Gold 120 C18, 5 µm, 250×4.6 mm from Dr. Maisch (Ammerbuch-Entringen, Germany). Total ionization chromatogram (TIC) for the schizokinen series (**4A**) and the extracted ion chromatogram (EIC) for each derivative (**4B-E**).

The obtained exact masses and their adducts are presented in **Fig 5A-D**. For the schizokinen series, we detected the following molecular ions and adducts; [M+H]^+^: m/z 387.1767, [M+H_2_O]^+^: m/z 404.1584 and [M+Na]^+^: m/z 409.1584 for N-deoxyschizokinen A (**5A**), [M+H]^+^: m/z 403.1845 and [M+Na]^+^: m/z 425.1651 for schizokinen A (**5B**), [M+H]^+^: m/z 405.2002 and [M+Na]^+^: m/z 427.1815 for N-deoxyschizokinen (**5C**), [M+H]^+^: m/z 421.1941 and [M+Na]^+^: m/z 443.1753 for schizokinen (**5D**).

**Figure 5.**
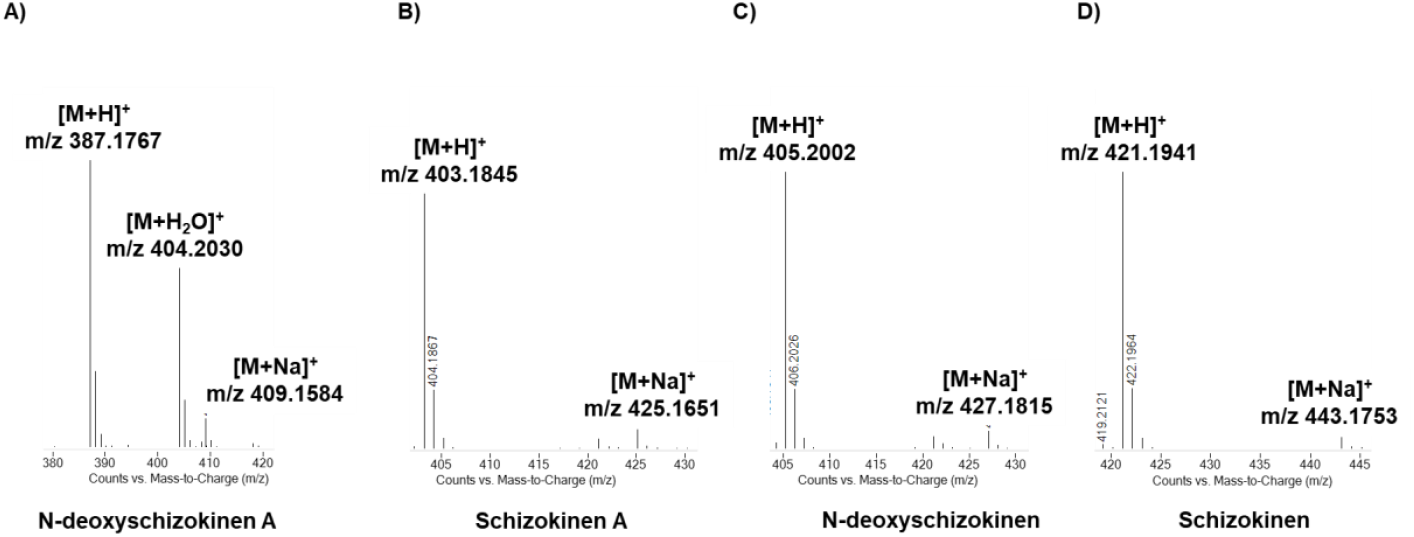
Obtained exact masses and their adducts for the schizokinen series. [M+H]^+^: m/z 387.1767, [M+H_2_O]^+^: m/z 404.1584 and [M+Na]^+^: m/z 409.1584 for N-deoxyschizokinen A (**5A**), [M+H]^+^: m/z 403.1845 and [M+Na]^+^: m/z 425.1651 for schizokinen A (**5B**), [M+H]^+^: m/z 405.2002 and [M+Na]^+^: m/z 427.1815 for N-deoxyschizokinen (**5C**), [M+H]^+^: m/z 421.1941 and [M+Na]^+^: m/z 443.1753 for schizokinen (**5D**).

### Tandem mass spectrometry and fragment assignment

The determination of the most characteristic fragments of the schizokinen series was completed through multiple fragmentation experiments, in which we determined the most recursive and intense daughter ions for each molecule, see **Table I** and **Figure 6**. To date, there was no complete set of information on the mass fingerprints of the schizokinen series. The only molecule which had information on its mass fragmentation pattern was schizokinen, published more than 10 years ago (Storey et al. 2006). As expected, the most characteristic fragments reported in the above-mentioned study were consistent with the ones we found:, m/z 133, m/z 243, m/z 361 and m/z 403. Furthermore, the collision energy used in the previous study (19-25 eV) was in a similar range than the one we found optimal (20 eV). Importantly, in our study, we also determined the daughter ions for three different collision energy experiments for the whole schizokinen series (**Table I**). Thus, we picked the most characteristic fragments for each molecule and postulated the fragmentation pathways, which can be observed in **Fig 6A-D**. To the best of our knowledge, the fragmentation patterns of schizokinen A, N-deoxy schizokinen and N-deoxy schizokinen A have not been reported to date.

**Table I.** Analytical features for the detection of the schizokinen series.

| Molecule | Schizokinen<br>$C_{16}H_{28}N_4O_9$ | N-deoxyschizokinen<br>$C_{16}H_{28}N_4O_8$ | Schizokinen A<br>$C_{16}H_{26}N_4O_8$ | N-deoxyschizokinen A<br>$C_{16}H_{26}N_4O_7$ |
| --- | --- | --- | --- | --- |
| Exact mass (Da) | 420.19 | 404.19 | 402.18 | 386.18 |
| Retention time column MN (min) | 7.05 | 5.73 | 8.41 | 17.4 |
| Retention time column DM (min) | 7.52 | 6.09 | 8.99 | 18.25 |
| Used collision energy (eV) | 10, 20, 30 | 10, 20, 30 | 10, 20, 30 | 10, 15, 20 |
| MS/MS fragments for 10eV (m/z) | 133, 243, 361, 403, 375 | 100, 117, 178, 345, 387 | 193, 343, 361, 385 | 105, 119, 281, 369 |
| MS/MS fragments for 15eV (m/z) | - | - | - | 105, 119* |
| MS/MS fragments for 20eV (m/z) | 116, 133, 201, 243, 261, 343, 361, 375, 403 | 56, 100, 117, 133, 178, 227, 243, 309, 327, 345, 387 | 56, 133, 178, 193, 211, 246, 343, 361, 385 | 77*, 105, 119* |
| MS/MS fragments for 30eV (m/z) | 56, 116, 133, 168, 183, 201, 243, 261, 325, 343, 361, 403 | 56, 91, 100, 117, 133, 168, 178, 193, 227, 317, 327, 345 | 56, 133, 178, 193, 211, 228, 246, 283, 343, 361 | - |
| Most characteristic fragments (m/z) | 133, 243, 361, 403 | 117, 227, , 345, 387 | 193, 343, 361, 385 | 105 |

**MN:** Nucleodur C18 Isis, Macherey-Nagel
**DM:** Reprosil Gold 120 C18, Dr. Maisch
\*: very low intensity

**Figure 6.**
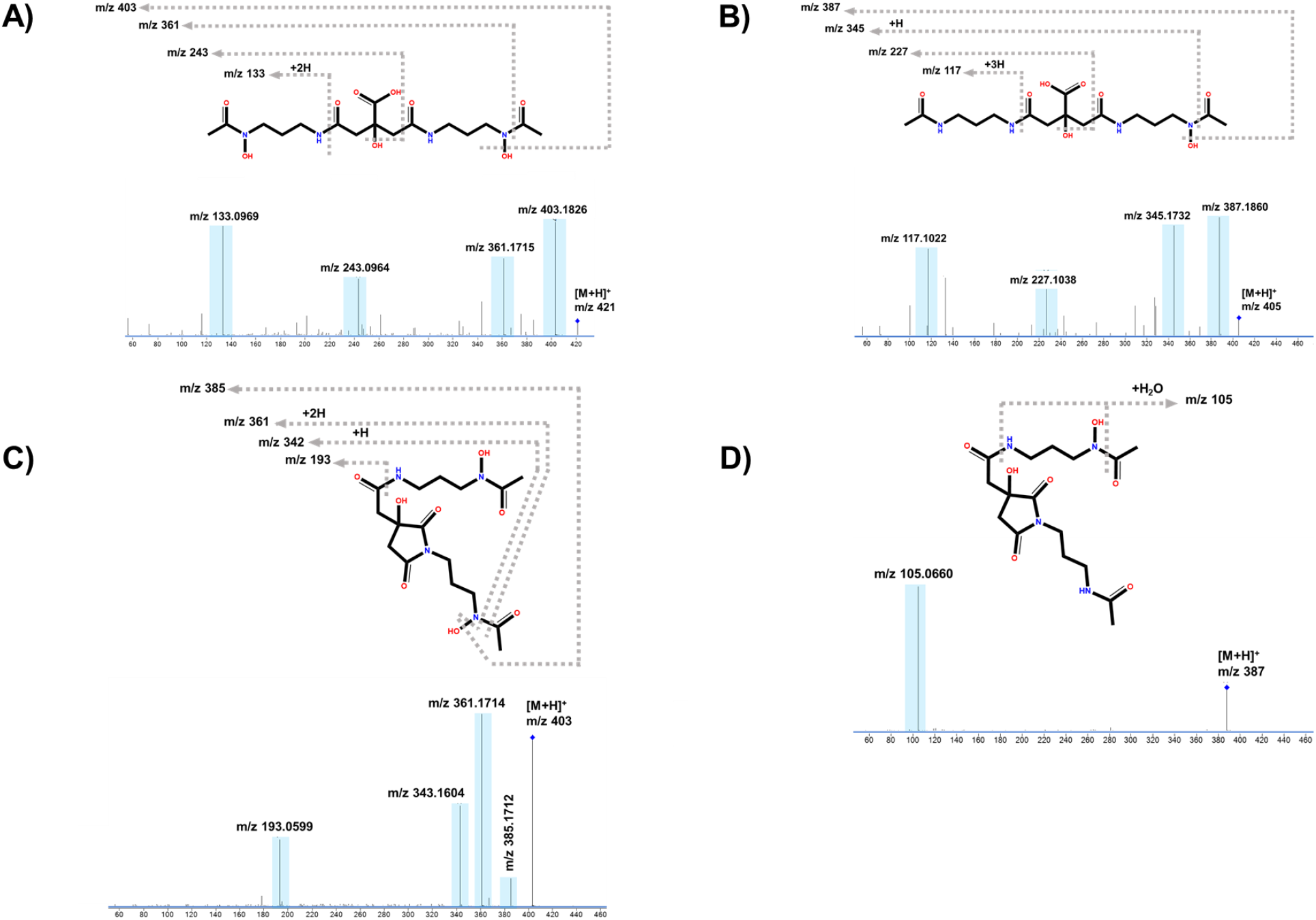
Mass tandem spectra and proposed fragmentation path for A) schizokinen, B) N-deoxyschizokinen, C) schizokinen A and D) N-deoxyschizokinen A.

The fragmentation of the all the schizokinen series occurred in a similar way in all the derivatives. There were four observed mechanisms for the fragmentation of the molecules, which were 1) cleavage of the amide bond, 2) cleavage of the hydroxamate bond, 3) alpha-cleavage and 4) neutral water losses. The reported fragmentation pathways are in agreement with reported collision-induced dissociation experiments performed in different groups of siderophores (Mawji et al. 2008).

Remarkably, N-deoxyschizokinen A showed a different behavior to its sibling molecules. Firstly, N-deoxyschizokinen A did not seem to fragment in the same fashion than the other schizokinen derivatives, since it produced a unique peak at m/z 105, which is not present in the other molecules. In addition, the intensity of this peak overwhelmed any other potential fragment being generated (**Fig 6D**). Furthermore, the analysis of the fragmentation of N-deoxy-schizokinen A indicated that its main fragment, m/z 105, corresponded to an adduct bearing a water molecule, which is not very often observed, but that has been reported (Housman et al. 2014). The elucidation of the fragment was determined by high resolution mass spectrometry, where the obtained exact mass, m/z 105.066, produced the molecular formula C_3_H_9_N_2_O_2_. The accuracy of the measurement was within 2 ppm, where the measured mass was m/z 105.0660 compared to the calculated m/z 105.0659. The determination of the degree of unsaturation (DU) of the fragment revealed a 0.5 DU, which is indicative of a radical. Finally, m/z 105 fragment was assigned to the cleavage of the longest arm of the molecules (1-amino-3*-*(*N*-hydroxy-*N*-acetyl)-aminopropane), where it is cleaved in both amide and hydroxamate bonds yielding a fragment with the neutral composition of 3-(hydroxy)propan-1-amine, in turn, this fragment associated with a water molecule to produce the unique and most intense peak observed in the fragmentation of N-deoxy-schizokinen A (**Fig 6D**). Finally, N-deoxyschizokinen A was a more delicate molecule for obtaining its fingerprint, since by increasing the collision energy in the same levels than its sibling molecules, the parent ion was decomposed entirely, and a single m/z 105 peak could be observed.

As these molecules are built up from almost the same starting material, it is reasonable to think that these molecules share common daughter ions after the mass tandem fragmentation. Therefore, during our experiments, we observed common fragments across 3 out of the 4 schizokinen derivatives, which were: m/z 56, m/z 73, m/z 100, m/z 133 and 193. It is important to remark that all but m/z 56 were chosen as MS2 identifiers for the schizokinen derivatives (see **Table I**). Despite m/z 56 was a good fingerprint for the detection and identification of the schizokinen derivatives, its low intensity across the samples made its selection difficult.

Finally, we wanted to determine if mass fragmentation platforms, such as CFM-ID (Allen et al. 2014), were able to correctly define the main fragments found in the schizokinen series. Surprisingly, the prediction made for the web server CFM-ID (supplementary info) was of good quality when compared with the empirical data (**Fig 6**). Although, we observed that the lower intensity ions were not successfully predicted, this could very well be a factor related with the instrument itself. Thus, data acquisition in different vendor instruments would be ideal to answer this question. A main disagreement was found for N-deoxyschizokinen A (MW: 386.18), which diverged of the expected fragmentation pathway. Thus, the empirical data for this molecule showed a main peak at m/z 105, which was not predicted by this platform. The m/z 105 corresponded to a water adduct, thus it is likely the reason of its absence from the prediction. Similarly, the core fragment (m/z 87) without the water molecule was not present in the prediction. Despite this observation, this platform is still useful to completement the experimental data and therefore, confirm and assign mass tandem fragments.

### 3.5 Molecular networking of the schizokinen series

The construction of the molecular networking of the schizokinen derivatives (**Fig 7**) provided a network with 17 nodes. The colored octagons represent the schizokinen derivatives, while the orange circles are unidentified analogues that may be a result of in-source fragmentation, stable decomposition by-products, as well as very low concentration derivatives. The molecular networking fingerprint for all the schizokinen derivatives, is useful for the identification of these metabolites in large datasets, which are complex to analyze manually, but that can be deduced by using metabolomic tools, such as the GNPS platform. Thus, we deposited the mass fragmentation data for the four schizokinen derivatives on a publicly available server to provide other researchers with this valuable information. The dataset can be retrieved from the Center for Computational Mass Spectrometry at the University of California, San Diego (massive.ucsd.edu) under the identification number MSV000088674.To conclude, molecular networking has just been implemented in a two-step native electrospray ionization-mass spectrometry method for the characterization of metal-binding compounds, such as siderophores. Thus, we think that by contributing our data can also help to speed up its characterization of siderophores (Aron et al. 2022).

**Figure 10.**
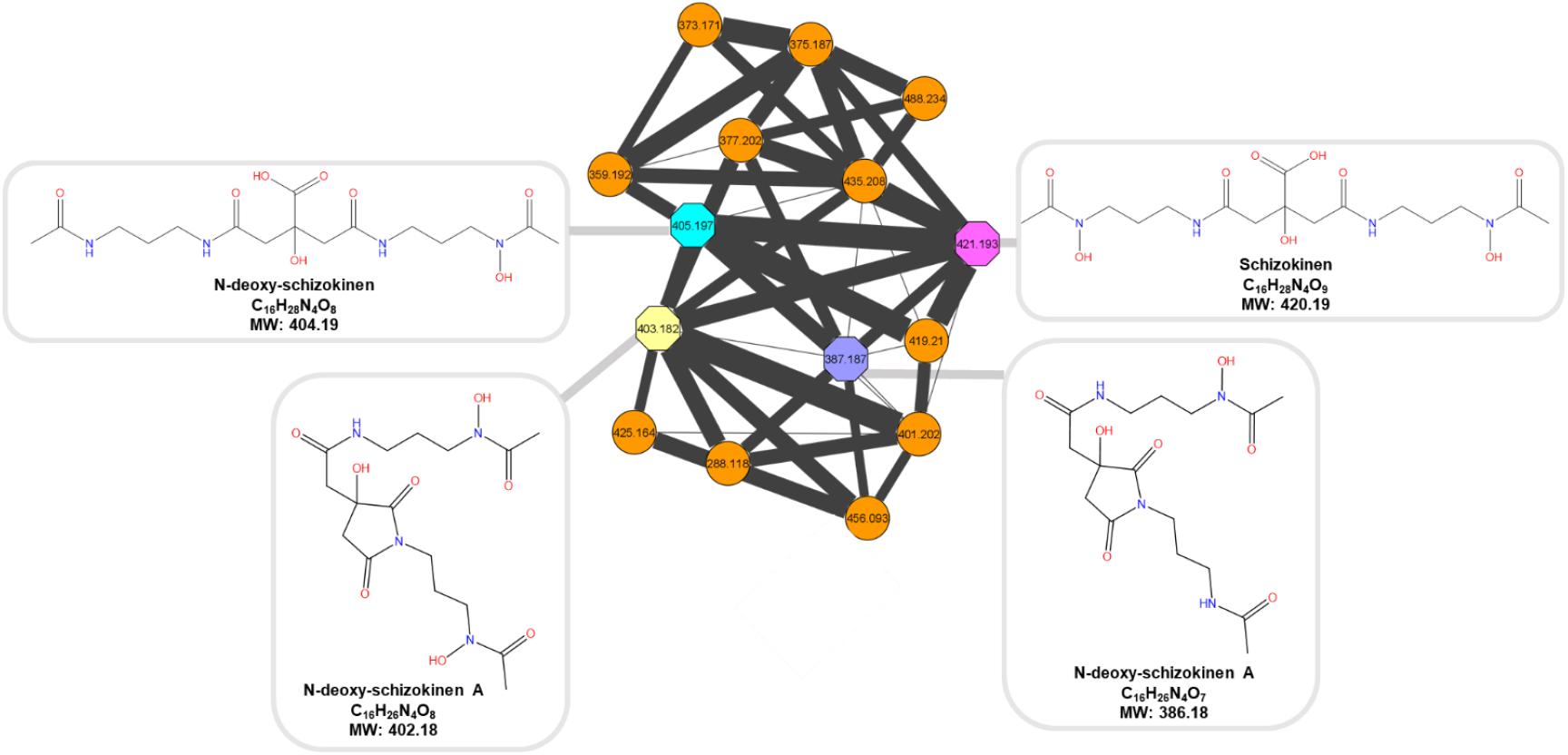
Molecular networking for the schizokinen series. Octagons represent the [M+H]^+^ of the schizokinen derivatives, while orange circles are the schizokinen low concentration analogues, as well as stable decomposition byproducts. Data obtained in positive mode.

## Conclusion

We established a specific methodology for the unambiguous identification in a single experiment of the siderophores schizokinen, schizokinen A, N-deoxy schizokinen and N-deoxy schizokinen A by using LCHRMS-MS. The present study offers four layers of confirmation; retention time (HPLC), exact mass (MS), mass fragmentation pattern (MS/MS) and the molecular networking fingerprint for the detection, identification, and potential quantification for each of these molecules. This work aims to improve the efficiency in the identification and characterization of the schizokinen siderophores series in different producing or containing sources, such as bacterial, algae and fungal cultures, in addition to clinical and soil samples.

## Supporting Information and Data Availability

Spectral data with different collision energies and the list of the CFM-ID predicted ions are provided in the supplementary information. The LCMS dataset can be retrieved from the Center for Computational Mass Spectrometry at the University of California, San Diego (massive.ucsd.edu) under the identification number MSV000088674.

## Acknowledgements

IS is thankful for the support provided by the Chemistry faculty of LMU and to the dean, Prof. Dr. Philip Tinnefeld.

